# Individual variation of the masticatory system dominates 3D skull shape in the herbivory-adapted marsupial wombats

**DOI:** 10.1101/692632

**Authors:** Vera Weisbecker, Thomas Guillerme, Cruise Speck, Emma Sherratt, Hyab Mehari Abraha, Alana C. Sharp, Claire E. Terhune, Simon Collins, Steve Johnston, Olga Panagiotopoulou

## Abstract

**Background:** Within-species skull shape variation of marsupial mammals is widely considered low and strongly size-dependent (allometric), possibly due to developmental constraints arising from the altricial birth of marsupials. However, species whose skulls are impacted by strong muscular stresses – particularly those produced through mastication of tough food items – may not display such intrinsic patterns very clearly because of the known plastic response of bone to muscle activity of the individual. In such cases, shape variation should not be dominated by allometry; ordination of shape in a geometric morphometric context through principal component analysis (PCA) should reveal main variation in areas under masticatory stress (incisor region/zygomatic arches/mandibular ramus); but this main variation should emerge from high individual variability and thus have low eigenvalues.

**Results:** We assessed the evidence for high individual variation through 3D geometric morphometric shape analysis of crania and mandibles of thre species of grazing-specialized wombats, whose diet of tough grasses puts considerable strain on their masticatory system. As expected, we found little allometry and low Principal Component 1 (PC1) eigenvalues within crania and mandibles of all three species. Also as expected, the main variation was in the muzzle, zygomatic arches, and masticatory muscle attachments of the mandibular ramus. We then implemented a new test to ask if the landmark variation reflected on PC1 was reflected in individuals with opposite PC1 scores and with opposite shapes in Procrustes space. This showed that correspondence between individual and ordinated shape variation was limited, indicating high levels of individual variability in the masticatory apparatus.

**Discussion:** Our results are inconsistent with hypotheses that skull shape variation within marsupial species reflects a constraint pattern. Rather, they support suggestions that individual plasticity can be an important determinant of within-species shape variation in marsupials (and possibly other mammals) with high masticatory stresses, making it difficult to understand the degree to which intrinsic constraint act on shape variation at the within-species level. We conclude that studies that link micro- and macroevolutionary patterns of shape variation might benefit from a focus on species with low-impact mastication, such as carnivorous or frugivorous species.

## Introduction

Much of mammalian diversity is reflected in the morphology of the skull (the cranium and mandible), which caters to the reception of sensory input, accommodates a large brain, and maintains a specialized masticatory apparatus [1]. However, mammalian skull diversity does not reflect a random walk through all possible adaptive shapes, but instead evolves within complex intrinsic constraints, which can be phylogenetic, developmental, and/or genetic [2–6].

Hypotheses of how intrinsic constraints shape mammalian skull diversity have been best articulated for marsupial mammals. Their unique altricial birth, and particularly the very early onset of a long suckling phase while the skull is barely developed, is thought to constrain the capacity for the marsupial skull to evolve [7–10]. This is supported by findings of lower cranial and mandibular shape disparity compared to placentals [7, 10–12]. Because the marsupial diversity constraint is thought to be ontogenetic, it is expected to limit the ability of individuals within a species to respond to selection [13, 14]. Several studies have indeed characterized marsupial skull shape over short evolutionary time scales (within species or between close relatives) and suggested that marsupial cranial shape variation mainly changes with size as an allometric “line of least resistance” [3, 4, 12, 15–18]. This is consistent with suggestions that developmental integration may manifest in strong developmental and static allometry [19], which has been specifically posited for marsupials due to their altricial birth [4].

Suggestions of a constraint on marsupial skull shape through static allometry are at odds with several geometric morphometric studies that show only low or moderate levels of static allometry in marsupial crania [15, 20–22]. Two recent studies on kangaroos even suggested that allometry plays a lesser role in shaping cranial variation in this group, instead positing fast adaptation or individual developmental plasticity of the masticatory apparatus as the main driver [23, 24]. This is consistent with suggestions that masticatory biomechanics may impact individual cranial shape to such a degree that developmental constraints either do not contribute much to within-species shape variation, or can be obscured by individual differences in mastication due to the bone’s plastic response to mastication stresses [6, 25–27]. Thus, high levels of biomechanical impact on the cranium may cause an important disjunct between within-species (static) *versus* evolutionary patterns of shape variation. This would add an important caveat to the co-interpretation of within- and between-species shape variation because they may have independent sources [13].

In this study, we test the “biomechanics hypothesis” through three-dimensional (3D) geometric morphometric analysis of cranio-mandibular shape variation in three species of specialized grazing marsupial, the wombats. Wombat skulls are under high masticatory stress due to a diet of fibrous grasses [28–31], making them an ideal test case for the “biomechanics hypothesis”. In particular, if masticatory stress is a driver of skull shape at the level of individuals, we should expect the main shape variation to occur in areas of high stress [the zygomatic arch, mandibular ramus, and the incisor alveolae; 28, 29-32] according to individual feeding habits. This variation should be dominated by the co-variance between cranium and mandible, be independent of size (*contra* hypotheses of an allometric “line of least resistance”). Lastly, if within- and between-species shape variation arises from different drivers, we expect that overall shape variation patterns should differ when comparing within-species and between-species variation.

For the case that individual feeding behaviour is a major determinant of within-species shape, it is also predicted that the main shape variation within our dataset should be visually similar in all species. However, this main variation should explain much of the overall variation as determined by conventional principal components analysis (PCA) [11, 33], with individuals showing different emphases on different parts of their masticatory system. However, assessing predictions of shape variation for specific shape partitions is challenging. This is because the main techniques for summarizing shape variation are ordination techniques such as PCA, which are based on the ordination of the landmark coordinate’s variance/covariance, rather than the range of the actual variation of the landmark positions. This procedure is blind to biological processes and prone to misleading conclusions [19, 34, 35], particularly where shape variation is visualized along ordination axes with low eigenvalues, as expected here. Thus, ordination-based interpretations of shape variation can lead to erroneous conclusions visually (e.g. variation is “smoothed” and shown without the context of other axes of variation [36]), procedurally (e.g. circularity when using PCA to both propose and test hypotheses), and statistically (e.g. when only a few axes are used for analyses or distinguishing groups or when the groups’ main axes of variation differ).

To produce a quantitative assessment of shape variation within our sample, beyond PCA, we here use a permutation test of the prediction that landmarks of Procrustes-superimposed mandibles and crania in areas under high masticatory stress should vary more than the overall landmark distribution in hypothetical PC1 shape extremes. In addition, if our prediction of high individual variation is correct, we predict that the magnitudes of shape variation along PC1 should not consistently match those of pairs of actual landmark configurations in extreme specimens 1) along the main axis of shape variation (PC1), and 2) at extreme ends of Procrustes space. Using this procedure, we show that the morphospace of wombat shape indeed behaves as predicted, with low allometry, small eigenvalues of the first principal component, and limited correspondence between PC1 shape variation and actual specimen shapes.

## Results

### Disparity comparison, allometry and covariation

The northern hairy-nosed wombat had lower cranial (but not mandibular) Procrustes variance than common and southern hairy-nosed wombats (Table 1). There was no significant effect of sex on either shape or centroid size in the two species with sex data available (Additional File 1).

**Table 1:**
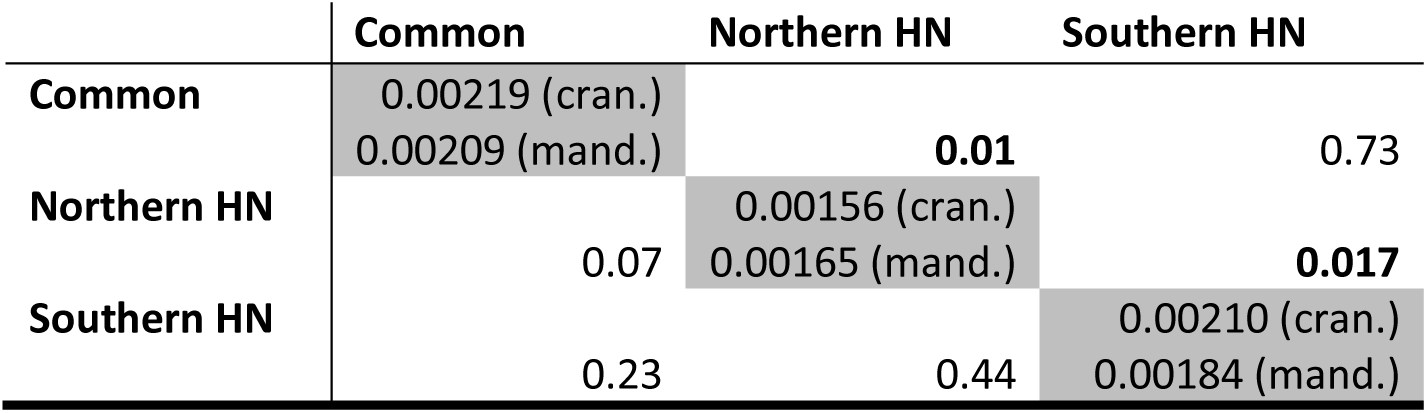
Procrustes variance comparisons among wombat species using pairwise comparison tests with 1000 replicates. Upper/lower diagonals: p-values for comparisons of crania (cran.)/mandibles (mand.). respectively. Procrustes variance values for each species are on the diagonal. HN=Hairy-nosed. Bold *p*-values are significant at *p*<0.05.

There was either no, or very little, allometry in the cranium and mandible within all wombat species (Table 2). A small but significant effect of allometry exists when the crania or mandibles of all species are analysed together (crania: R^2^=0.08; F=5.65; *p=*0.004; mandibles: R^2^=0.06; F=3.30; *p=*0.007); plotting cranial and mandible shape against centroid size (Additional File 2) revealed that this is driven by the different shape of the smaller southern hairy-nosed wombats, whereas northern hairy-nosed and common wombats overlap according to size and shape.

**Table 2:**
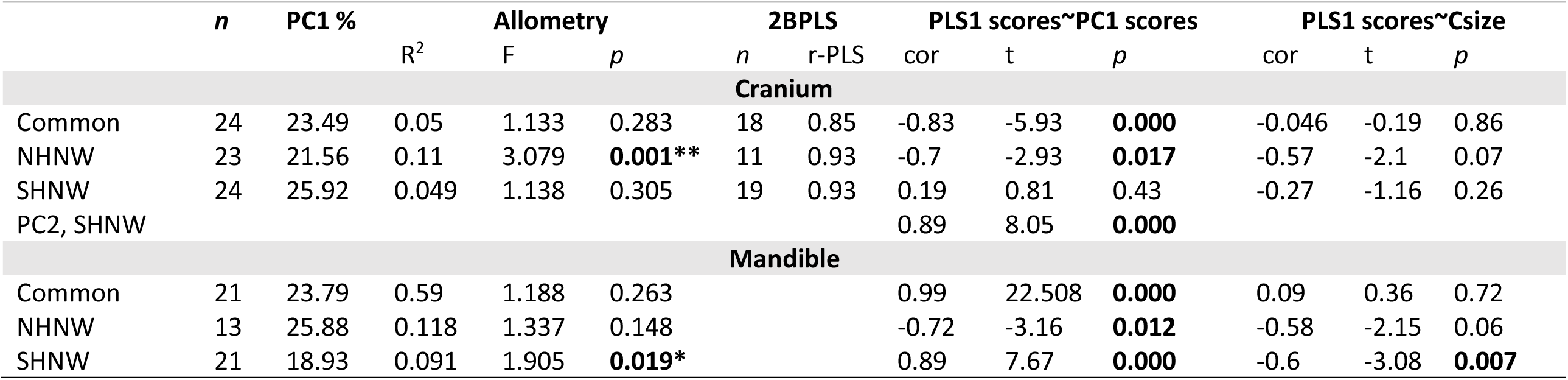
Summaries of the PCA analyses, Allometry analyses, two-block partial least squares (2BPLS) analyses, and correlation statistics between and PLS 1 scores with the species’ PC1 scores and centroid sizes. *n*, number of specimens used in the analyses; R^2^, coefficient of determination of linear models; t, t-statistic of correlation test; *p*: p-value, CW = common wombat; NHNW = northern hairy-nosed wombat; SHNW= southern hairy-nosed wombat

| | *n* | PC1 % | $R^2$ | Allometry | | 2BPLS | | PLS1 scores~PC1 scores | | | PLS1 scores~Csize | | |
| --- | --- | --- | --- | --- | --- | --- | --- | --- | --- | --- | --- | --- | --- |
|  |  |  |  | F | *p* | *n* | r-PLS | cor | *t* | *p* | cor | *t* | *p* |
| **Cranium** |  |  |  |  |  |  |  |  |  |  |  |  |  |
| Common | 24 | 23.49 | 0.05 | 1.133 | 0.283 | 18 | 0.85 | -0.83 | -5.93 | **0.000** | -0.046 | -0.19 | 0.86 |
| NHNW | 23 | 21.56 | 0.11 | 3.079 | **0.001\*\*** | 11 | 0.93 | -0.7 | -2.93 | **0.017** | -0.57 | -2.1 | 0.07 |
| SHNW | 24 | 25.92 | 0.049 | 1.138 | 0.305 | 19 | 0.93 | 0.19 | 0.81 | 0.43 | -0.27 | -1.16 | 0.26 |
| PC2, SHNW |  |  |  |  |  |  |  | 0.89 | 8.05 | **0.000** |  |  |  |
| **Mandible** |  |  |  |  |  |  |  |  |  |  |  |  |  |
| Common | 21 | 23.79 | 0.59 | 1.188 | 0.263 |  |  | 0.99 | 22.508 | **0.000** | 0.09 | 0.36 | 0.72 |
| NHNW | 13 | 25.88 | 0.118 | 1.337 | 0.148 |  |  | -0.72 | -3.16 | **0.012** | -0.58 | -2.15 | 0.06 |
| SHNW | 21 | 18.93 | 0.091 | 1.905 | **0.019\*** |  |  | 0.89 | 7.67 | **0.000** | -0.6 | -3.08 | **0.007** |

The two-block partial least squares analyses revealed high correlation coefficients between crania and mandibles (r-PLS) around 0.90 (Table 1). 2BPLS scores were highly correlated with PC1 scores in all cases except for the cranium of southern hairy-nosed wombats, where instead PC2 and the 2BPLS scores were strongly correlated. By contrast, 2BPLS scores were only correlated with centroid size in southern hairy-nosed wombat mandibles.

Replicating the analyses reported in Table 1 and 2 without surface semilandmarks and with only fixed semilandmarks revealed very few differences. Using fixed and curve semilandmarks only did not change the significance levels of any analysis, and using fixed landmarks changed the significance levels only in cases where significance levels were already close to the p = 0.05 cut-off (Additional File 3)

### PCA exploration and landmark position variation

PC1 explained little within-species shape variation in all species (18% to 25% Table 2). PCA of all wombats separated cranial and mandibular shapes of the two wombat genera on the first principal component (PC1; accounting for 45% and 30% of cranial and mandibular variation, respectively; Figure 1). PC1 of the sample of *Lasiorhinus* (accounting for 40% and 30% of cranial/ mandibular variation, respectively), described the differences between northern and southern hairy-nosed wombats with some very minor overlap (Figure 1).

**Figure 1:**
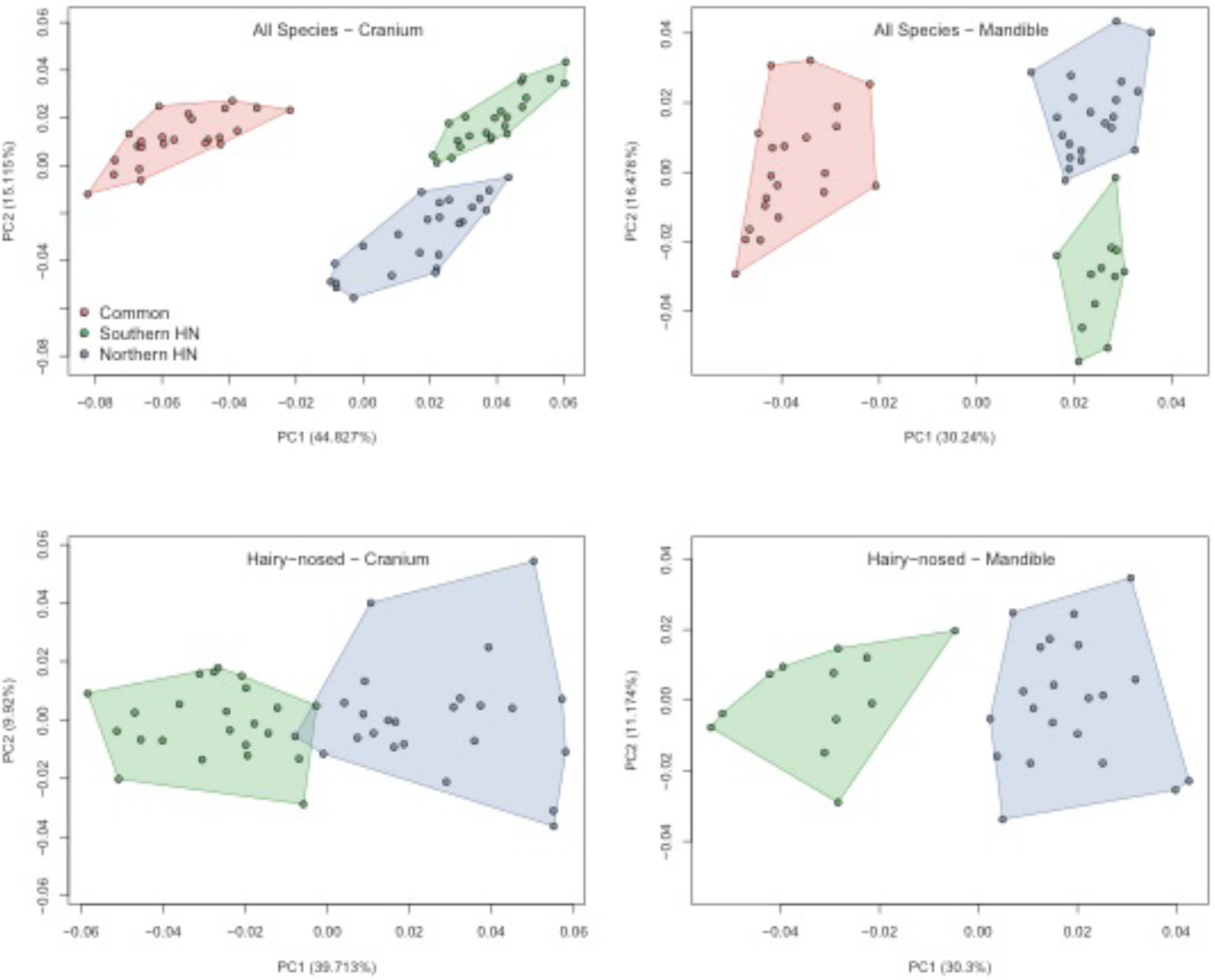
Principal Component 1 vs. 2 plots of cranial (left) and mandibular (right) shapes, showing distributions of specimens in the all-wombat (above) and hairy-nosed wombat (below) sample.

Heat plots of landmark displacement between extreme PC1 shape configurations within species (Figure 2) confirm our expectation that the overall main variation relates to the muzzle, zygomatic arches, mandibular ramus at the masticatory muscle attachments (masseteric fossa, coronoid process, and angular shelf) and incisor alveolae. In addition, in all three species, longer, more ventral rostral regions coincide with less flared and more ventrally placed zygomatic arches. PC1 heatplots of all wombats and the hairy-nosed wombats only (Figure 3) – describing the differences between the two genera and hairy-nosed wombat species, respectively - show limited similarity to the within-species patterns, with the zygomatic arch or mandibular muscle attachments not showing much displacement between PC1 extremes. PC1 variation of hairy-nosed wombats differed from the within-species variation by reflecting extensive displacement in the occipital region and variation in the frontal region. In the mandible, the symphysis did not vary among hairy-nosed wombat species; the distal edge of the angular process varied most between the hairy-nosed wombats, together with a change in angle of the mandibular condyle. Relative to hairy nosed wombats, common wombats have a more ventrally directed masseteric scar, anteriorly elongate nasal region, extensive ventral displacement of the premaxillary suture, and a more dorsally placed palate. Further, the cranial and mandibular molar rows are more dorsally situated, and the mandibular condyle is more medially directed.

**Figure 2:**
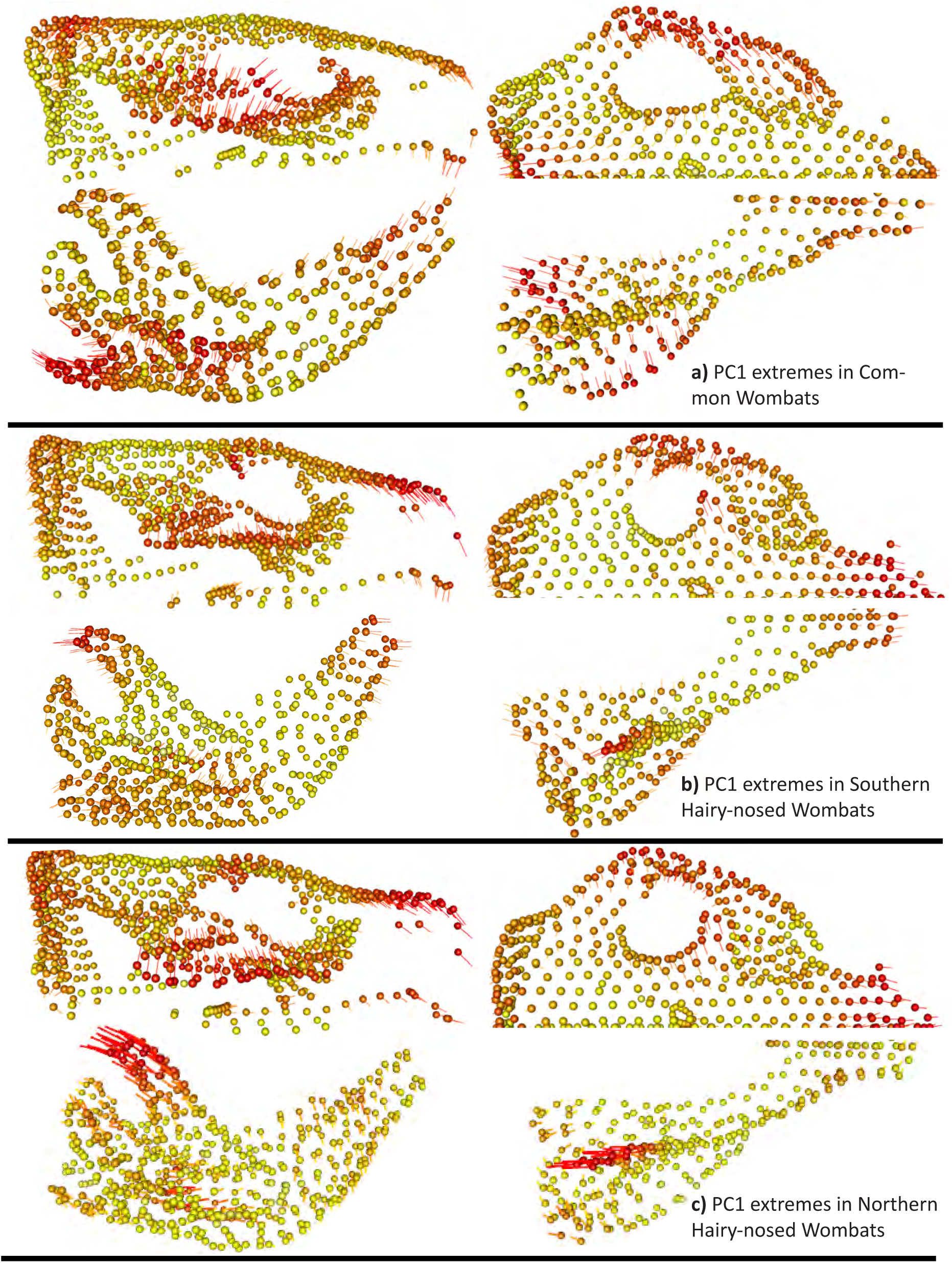
Heat plots representing the difference in landmark position between the two most extreme specimens along PC1. Dots are the position of one landmark of one PC extreme, lines represent the displacement of the same landmark in the other. Colour heat reflects displacement magnitude (red/yellow = high/low displacement).

**Figure 3:**
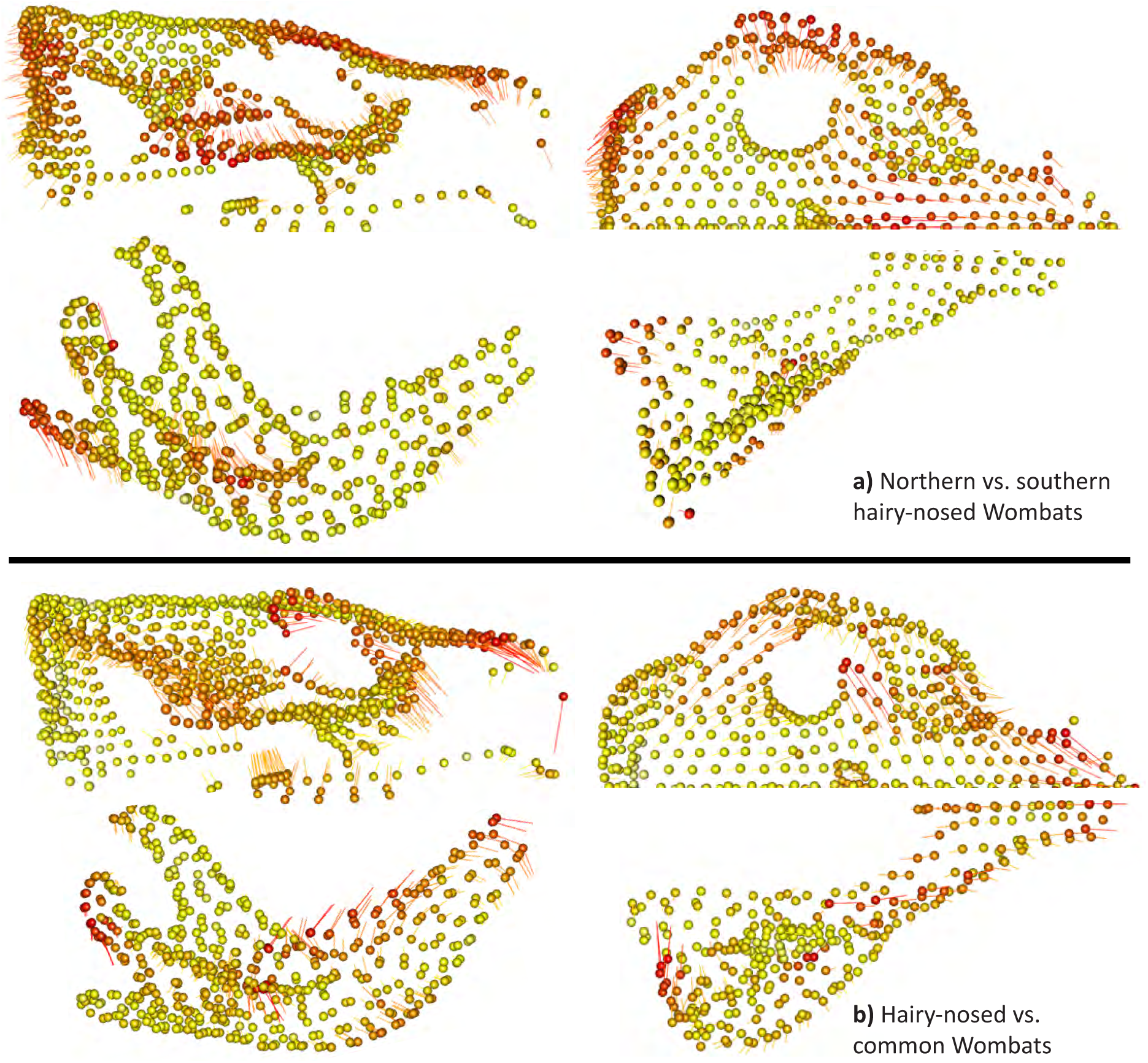
Heat plots representing the range of landmark displacement along PC1 in a) the all-wombat sample and b) in the sample of all hairy-nosed wombats, grouping the species of the genus.

### Landmark position variation test

The results of our landmark variation tests are summarized in Figure 4, and the corresponding specimen configurations for each within-species comparison at 100% confidence intervals are in Additional Files 4 and 5. Few results were robust to rarefaction to minimum partition sizes (black frames in Figure 4), so we cannot exclude that the significances might be an artefact of the larger numbers of landmarks in the other partitions. The best supported hypothesis across all comparisons was that the “remainder of cranium” partitions should vary less among all landmarks. Our hypotheses that the zygomatic arch, mandibular muscle attachments, and incisor area show greater landmark displacement were only partially supported, with highest support overall in comparisons of hypothetical PC1 configurations. Comparisons of specimen configurations on PC1 or Procrustes space extremes provided less support for our hypotheses, either by showing no significant displacement or by being significantly displaced but without being significantly different from the overall distribution of landmark displacements. Some comparisons between individual configurations even contradicted the visual impression from the heat plots by yielding lower displacements in the tip of rostrum/zygomatic arch partitions (yellow frames in Figure 3, see also Additional figures 4 and 5). This was particularly obvious in the whole-Procrustes space comparisons, showing that the variation according to PC1 is not a main component of shape across all of Procrustes space. Interestingly, all three hypotheses of cranial partition displacement were supported in comparisons of mean configurations of common and hairy-nosed wombats, while none was supported in mean species configurations between northern and southern hairy-nosed wombats.

**Figure 4:**
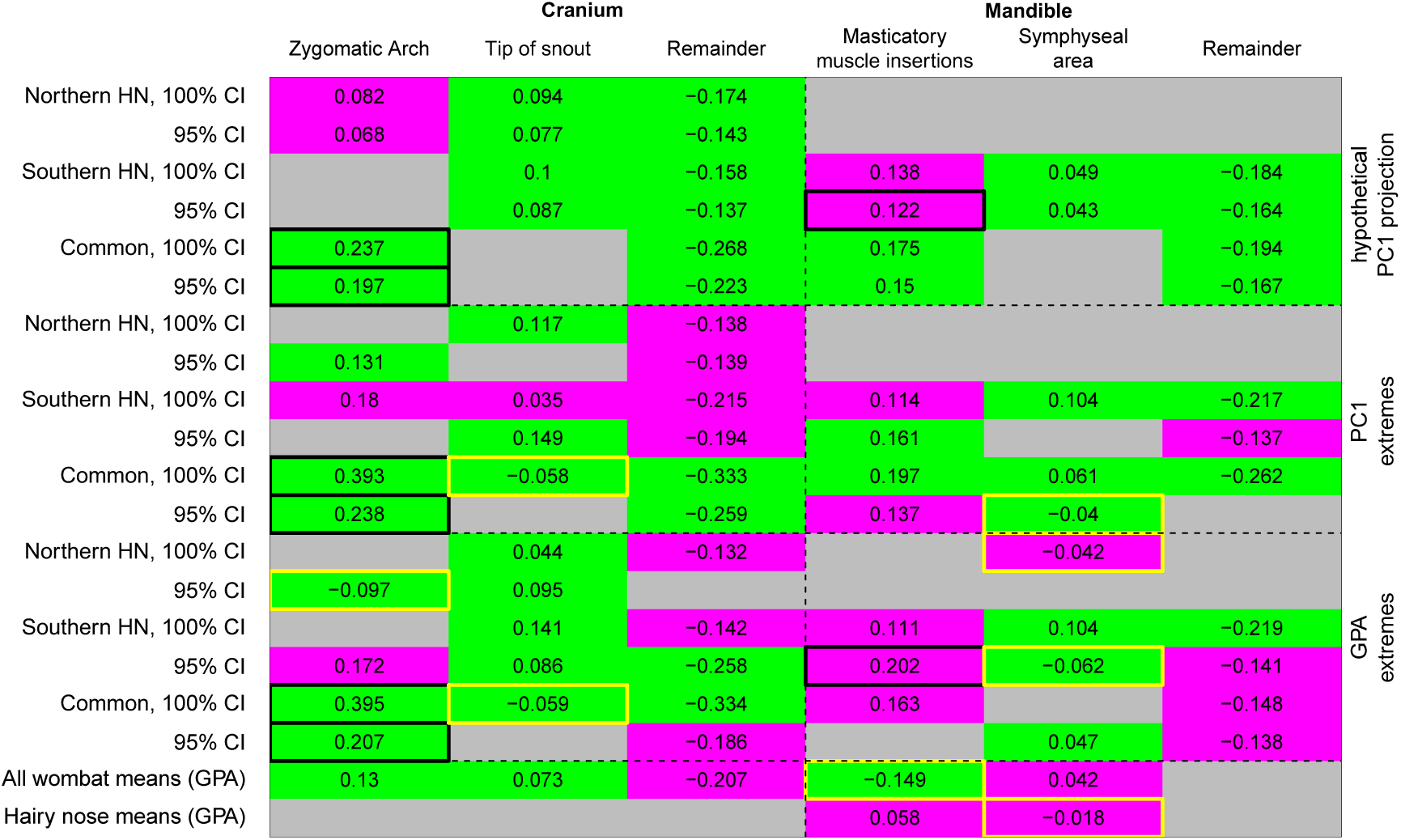
Landmark variation test results, between the most extreme specimens (100% and 95% CI) based on the hypothetical PC1 projection, PC1 and the GPA (rows). Negative and positive values: smaller and larger range of landmark variation than the whole cranium or mandible. Grey: no significant differences of landmark displacement magnitude (*p* > 0.001); Magenta: significant displacement differences undistinguishable from the full statistical distribution of landmarks (Bhattacharrya Coefficient *p* > 0.001). Green: significant differences in magnitude and statistical distribution. Black frames, significant also when rarefied. Yellow frames, results opposite to hypothesized effect.

**Figure 5:**
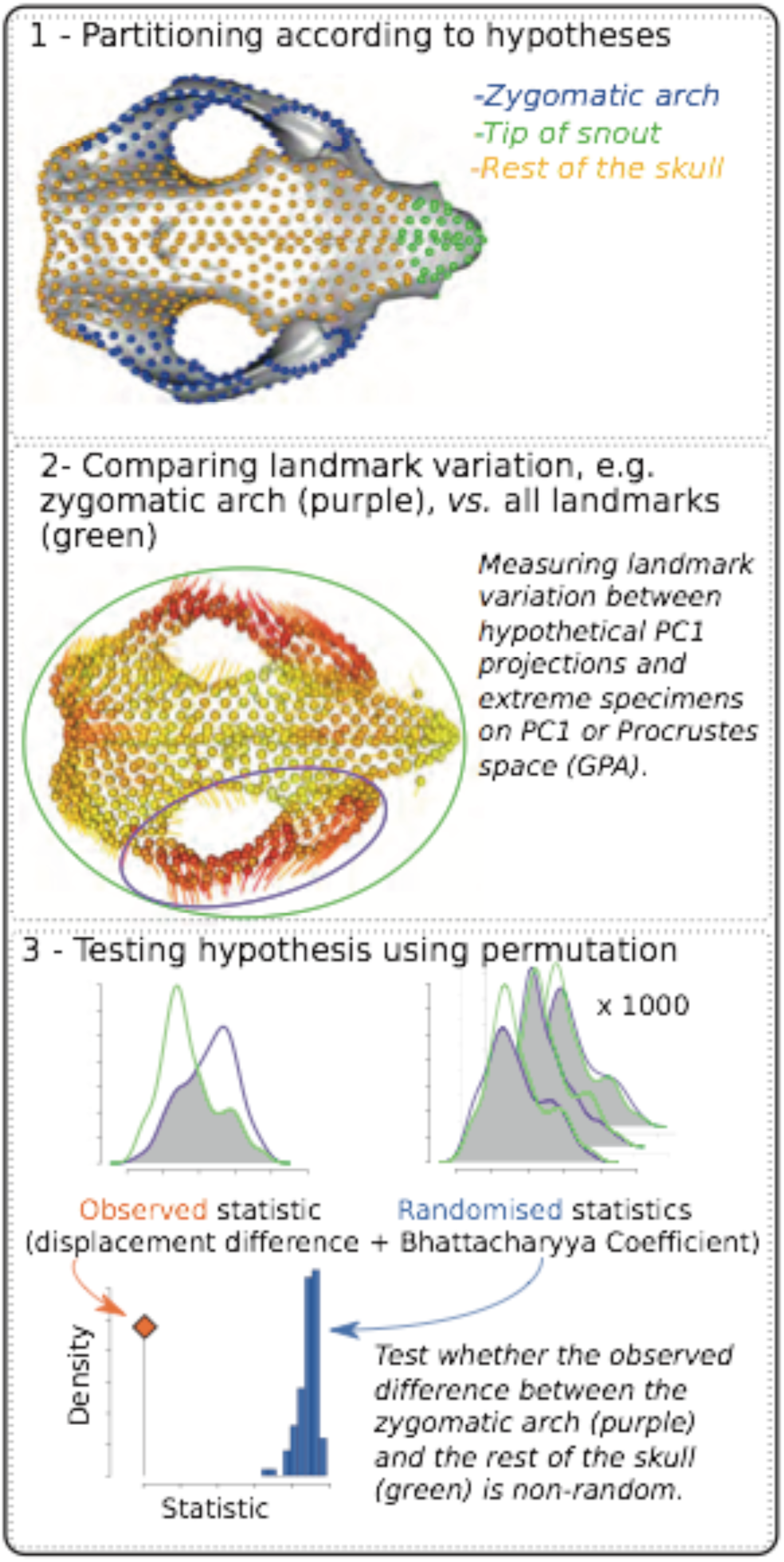
Outline of the landmark variation test. 1-identifying landmark partitions and statistical null. 2 – measuring distance between corresponding landmarks in pairs of specimen (visualized as lines) 3 - Permutation test to assess whether a partition varies more than the whole of the skull based on displacement difference and Bhattacharyya Coefficient.

## Discussion

Our results support our “biomechanics hypothesis” of a non-allometric, mastication-related driver of cranio-mandibular shape variation within wombat species. As expected for a skull shaped by masticatory biomechanics [29, 31, 32, 47], the heat plots of hypothetical PC1 shape extremes within species reveal strikingly uniform patterns of high landmark displacements in the zygomatic arches and rostra, also consistent with our hypothesis that these areas should vary most within species (in the mandible, displacement directions are not as uniform but still all occur in the muscle attachement sites). There is little indication that allometry plays any role in shaping within-species shape variation, with both overall shape and cranio-mandibular shape co-variation not related to centroid size (and shape co-variation instead strongly correlated with the main ordinated variation).

We also find substantial support for our prediction of high levels of individual variation within each species, as expected if individual diet or feeding habit were a main driver of skull shape. In particular, the main axes of shape variation had low eigenvalues, with PC1 explaining at most 25% of variation. Our “landmark test” of landmark displacement magnitudes confirmed that these low eigenvalues are related to inconsistencies between ordinated and individual configurations. Specifically, our hypotheses of greater displacement in the “tip of rostrum” and “zygomatic arch” partitions are best-supported in the hypothetical configurations of PC1 extremes, with less support or even contradictory evidence in actual specimens with opposite PC1 scores or on opposite ends of Procrustes space. In addition, comparisons of actual specimens between PC1 extremes and in extremes of Procrustes space showed that displacement directions (which were not measured by our landmark test) in the areas under high displacement differed substantially in the actual specimens, particularly in the mandibles. Thus, despite their overall similarity, shape variation along PC1 within species clearly arises from individual shapes in areas of high mechanic stress that only reflect parts of the variation identified by PC1. This contrasts with predictions of individual shapes falling along a spectrum of change along a high-eigenvalue PC1, as expected under a scenario of constraint. Rather, it concurs with our expectation that within-species morphospaces can contain too much “noisy” variation to infer a constraint or interpret in a context of macroevolutionary patterns [6, 13, 27, 48].

While individual variability in our sample is high, shape variation within species tends to occur most frequently in areas of high masticatory stress, which also seems to cause the uniform within-species PC1 landmark displacement patterns. Remodelling of these areas according to individual mastication habit or dietary preferences thus seems a plausible source of individual variation. This is consistent with finds that the zygomatic arch grows differentially upon the onset of feeding in two marsupial species [Virginia opossums and New Guinea Quolls; 49, 50], and remodels under mechanical stress in mammals [26, 47, 51, 52]. A similar remodelling process may also cause the high shape variation in the occipital crest along our within-species PC1, which is under mechanical stress from the temporalis [31] and nuchal muscles [53]. Lastly, the genetically and spatially highly restricted population of northern hairy-nosed wombat showed only slightly lower cranial shape disparity, and similar overall landmark displacement patterns, compared to the other two wombat species; this again suggests a strong role of non-heritable feeding behaviours, rather than a role of genetic diversity [3], in shaping individual wombat skulls.

Our findings support previous concerns about allocating the contributions of individual variability and heritable adaptation in closely related mammals [6, 27]. In particular, the within-species variation occurs in parts of the skull that also adapt to a herbivorous lifestyle in macroevolutionary contexts [54–57], posing potential issues for distinguishing between individual plasticity and between-species differences across evolutionary time scales. For example, the within-species PC1 displacement patterns of cranial shapes resembles some aspects of landmark displacement between the mean of *Vombatus* and *Lasiorhinus*. However, this difference between genera has evolved over the last seven million years and likely reflects the larger masseter muscle [30, 31], and different angle of incision [58] of common wombats.

The ambiguity of distinguishing within- and between-species variation in closely related species might also play a role in recent debates surrounding the existence of a possible universal Cranial Rule of Evolutionary Allometry (CREA). This posits that closely related mammals tend to have longer rostra and narrower zygomatic arches as they increase in size [59], in a pattern that is highly similar to the one we found within wombats but without allometry. CREA was suggested for kangaroo crania [18], but contested in a slightly different sample and landmarking protocol of kangaroos [23, 24]. Instead, the authors postulated that a CREA-like pattern is not allometric and instead purely due to biomechanics. CREA-like shape variation is also found in other kangaroo species, and is allometric in tammar wallabies [60] and quokkas [20], but only partially so among the rock wallabies [15]. By contrast, geometric morphometric analyses of two marsupial species with a comparatively soft diet – the insectivorous *Dromiciops gliroides* and two species of the omnivorous *Caluromys* – show little cranial variation of zygomatic arch or rostrum shape, but strong-species allometry [21, 22]. The variable association of the CREA shape variation with allometry may therefore be related to varying biomechanical stresses on individual skulls depending on a species’ feeding ecology (but note that CREA has been shown within mongooses and fruit bats, which both do not feed on hard items [59]). Feeding ecology, or other biomechanical skull function, therefore seems to be an important consideration in the choice of species for intraspecific assessments of developmental constraints [see also 61].

In methodological terms, use of the landmark variation test presents a useful additional tool to the geometric morphometric toolkit for interpreting shape variation. In our specific case, the spread of variation across several low-eigenvalue ordination axes rendered visualization through scatterplots of two or three axes ambiguous (i.e. the space containing important information was more multidimensional); co-investigation of individual specimen configurations in Procrustes space improved our interpretation of the morphospace. Similar procedures can also support the interpretation of other analyses (e.g. understanding the Procrustes space variation underlying patterns of allometry or other variables). Note that, like all procedures that interpret Procrustes-based landmark variation, the landmark variation test can be subject to artificial signals of directional variation due to the least-squares fitting of Procrustes superimposition [62, 63]; this can be alleviated by using high numbers and even coverage of landmarks. Note that in the case of high landmark numbers, other analyses may need to be assessed for robustness towards high numbers of landmarks [42]. Our approach also has the caveat that permutation tests have low resolution on small distributions [64], such as the occipital crest in our wombats. In such cases, conventional visual assessments of landmark displacement might be a better, if less quantitatively tractable, approach.

## Conclusions

Our results point towards a mostly biomechanically caused, mastication-related drive of wombat skull shape variation, suggesting that important drivers of shape macroevolution, such as a possible constraint on marsupial skull shape, may only emerge above the species level in some mammals [6, 13, 27, 48]. This posits an important challenge in testing hypotheses of constraints, and identifying differences between heritable and epigenetic variation, within mammalian skulls [13]; this is already being acknowledged, for example in studies that account for population-level shape “noise” in tree dating [65]. More detailed ontogenetic studies of shape might allow the separation of “original” skull shape prior to the onset of feeding [as in 49, 50], might help separate epigenetic from developmental shape variation.

## Methods

### Data collection and landmarking

We collected 3D data from crania and mandibles of adult common wombats (*Vombatus ursinus*; 24 crania/21 mandibles), southern hairy-nosed wombats (*Lasiorhinus latifrons*; 24 crania/21 mandibles) and northern hairy-nosed wombats (*Lasiorhinus krefftii*; 23 crania/13 mandibles). Data were acquired using two computed tomography (CT) scanners (Toshiba Activion16 at the School of Veterinary Sciences, Gatton, The University of Queensland) and a Somatom Definition AS+ scanner at I-MED Radiology, Armidale). Scans were converted to 3D surface meshes in Mimics v.19 (Materialise, Belgium). Author CS repaired surface meshes of five specimens that had minor damage (Additional Table 3). All original scans and 3D data are available on MorphoSource (www.morphosource.com, Project P418). For a list of specimens, their provenance, animal ethics and permit numbers for all non-museum specimens, and repairs, refer to Additional File 6; for scan details, refer to MorphoSource. The disparate numbers of crania and mandibles derive from the fact that not all specimens had both crania and mandibles available, so that we included just crania or just mandibles to maximize sample sizes. Samples of common and southern hairy-nosed wombat were across their geographical range (respectively, east to west South Australia / from Tasmania to Queensland). We only had specimens of Northern hairy-nosed wombat samples from the single surviving population (Additional File 6). To test whether this resulted in lower shape variance, we employed Procrustes variance comparisons among the species. We also tested if sexual dimorphism of shape was a possible confounding factor in the two wombat species with sex data (southern and common wombat), by performing Procrustes ANOVAs of shape/size against sex.

Landmarks were digitized by CS in Viewbox 4 (dHAL). This involved placing 65 landmarks and 761 semilandmarks (261 curve and 500 surface) on the cranium and 35 landmarks and 542 semilandmarks (142 curve and 400 surface) on the mandible (Additional File 7, Additional File 8). Landmarks and curve semilandmarks were placed manually, and their position was used to automatically place the surface semilandmarks, based on a template of an undamaged specimen of southern hairy-nosed wombat (CLVSJR5). Curve and surface semilandmarks were slid over the 3D surface of each scan (as opposed to being interpolated without surface information) according to the bending energy criterion [37, 38] in Viewbox v. 4.0.1.7. Landmark coordinates were analysed in R [39] using geomorph v. 3.1.0 [40] for standard analyses and a new package, landvR (https://github.com/TGuillerme/landvR) [41] for the permutation tests. Coordinates were scaled and superimposed using generalized Procrustes analysis (GPA) without a sliding step (as sliding was done over the 3D geometries in Viewbox), and ordinated through PCA. Separate GPAs and PCAs were done for crania and mandibles in all specimens, for hairy-nosed wombats only, and separately for each species. Data and code for all analyses are available and repeatable as R markdown files at https://github.com/TGuillerme/landmark-test.

### Landmark sampling

Our landmarking protocol was designed to densely capture featureless areas of the skull, such as the cranial vault, forehead, or mandibular ramus. Although we used Viewbox to ensure that landmarks were slid over the actual 3D surface, rather than interpolated from just fixed landmarks/curve semilandmarks, the dependence of the semilandmarks of fixed landmarks raises the issue of pseudo-replication [34, 35]. However, for our landmark test, high numbers of fixed landmarks were important. To ensure our other analyses were not affected by the high number of semilandmarks [e.g. 42], we therefore re-ran all analyses with just fixed and curve semilandmarks, and just fixed landmarks.

### Allometry and covariation

To test our prediction that allometry should not be a main driver of within-species cranio-mandibular shape, we first performed a Procrustes ANOVA of shape predicted by centroid size. To assess how cranio-mandibular integration related to allometry and the main ordinated variation, we also conducted a two-block partial least-squares (2BPLS) analysis of cranio-mandibular covariation in specimens with crania and mandibles. This finds the axis of greatest shape covariation between the cranium and mandible and identifies how much of shape variation in each dataset is due to the covariation with the other. We then asked if cranio-mandibular co-variation was a likely driver of the main shape variation by correlating 2BPLS scores with PC1 scores within each species. We also asked if there was evidence of an allometric constraint on the co-variation, by correlating the 2BPLS scores with centroid size.

### Landmark variation test

We performed a permutation-based landmark position variation test to test two hypotheses of shape variation, 1) that areas of the skull under high masticatory stresses show the greatest variation magnitudes within species and 2) that patterns of landmark displacement differ within and among species. This procedure is sketched in Figure 4, detailed in Additional File 9 and implemented in the landvR package (including tutorial vignettes) accessible at https://github.com/TGuillerme/landvR [41]

### Partition selection

Based on the literature on wombat skull biomechanics [31–34, 41], we selected four partitions expected to vary most and one partition expected to not vary (as a null hypothesis). In the cranium, partitions expected to vary were 1) the zygomatic arch (262 landmarks) and 2) the “rostrum” partition (the incisor alveolae, premaxillary suture, and nasal bones immediately dorsal to this region; 66 landmarks). In the mandible, these were the mandibular “muscle attachment” partition (the masseter/pterygoid/temporalis attachments in the mandibular ramus; 142 landmarks) and the “anterior symphysis” partition (anterior to the molars at the incisor roots; 35 landmarks). Partitions not expected to vary were the rest of the cranium (501 landmarks) or mandible (400 landmarks) (Additional File 10). To account for different landmark numbers per partition, we also rarefied each partition to the minimum landmark number (66/cranium and 35/mandible).

### Comparison between pairs of specimens and mean shapes

We sampled specimen pairs to reflect within-species variance by selecting: 1) the projection of the hypothetical extremes of configurations along PC1; 2) landmark configurations of specimens with the highest *vs.* lowest PC1 score; and 3) the most different specimens in Procrustes space (based on distance from the mean shape – see Additional text 1). To increase reliability of our results, we also tested pairs of specimens within the 95% CI of each distribution. Additionally, we tested if differences in mean shapes among species correspond to our hypotheses of shape change.

### Testing differences between extreme specimens by partitions

For each specimen pair, we tested whether a partition exhibited greater displacement of landmarks compared to the rest of the cranium or mandible. For each pair, we 1) calculated the Euclidean distance between corresponding landmarks in both specimens in Procrustes space; 2) calculated the displacement difference (i.e. total landmark distance between two specimens) and the Bhattacharyya Coefficient [43–45] (i.e. overlap between the distance distributions – see Additional text 1); and 3) performed a permutation test [46] on both statistics by comparing them to the statistics of 1000 same-sized random partitions (Figure 5). As the great number of tests (96) will lead to a type I error inflation, we reduced our p-value acceptance threshold from the traditional 0.05 to 0.001 (0.1%)

## Declarations

### Ethics approval and consent to participate

Where specimens were not sourced from museum collections, they were sourced with ethics permissions as outlined in Additional Table 3.

### Consent for publication

Not applicable

### Availability of data and material

The raw CT scans for all specimens used, and their derivative 3D surface files used for landmarking, are available on MorphoSource (www.morphosource.com, Project P418). The coordinate data and code to repeat all analyses for this study is available as rmd files at https://github.com/TGuillerme/landmark-test. Our landvR package for implementation of the landmark variation test, including a step-by step vignette, is accessible on https://github.com/TGuillerme/landvR.

## Author Contributions

VW and OP designed the study, co-developed the landmarking protocol, and wrote the manuscript. VW wrote parts of the code. TG wrote the heatplot and landvR package, supported the writing of the remaining code and co-wrote the manuscript. CS developed the landmarking protocol, placed the landmarks and co-analysed data. ES provided statistical support and co-wrote the manuscript. ACS and CT co-wrote the manuscript and provided biomechanical advice; ACS also collected parts of the CT data. HMA, SNC, OP and SJ collected and segmented the 3D data.

## Acknowledgements

We are grateful to Kate Garland, Rebecca Morrison, Manuel Wailan and Catherine Mullins for technical support during the 3D reconstruction. We thank David Stemmer (South Australian Museum), Heather Janetzki (Queensland Museum), and Alan Horsup (Queensland Department for Environment and Science) for access to wombat skulls. Thanks also to the staff at the Small Animal Clinic at the University of Queensland, and at I-med Radiology, Armidale, for allowing the scanning of specimens. Thank to Suren Rathnayake (University of Queensland) for insightful comments on the landmark variation test.

